# UK B.1.1.7 variant exhibits increased respiratory replication and shedding in nonhuman primates

**DOI:** 10.1101/2021.06.11.448134

**Authors:** K. Rosenke, F. Feldmann, A. Okumura, F. Hansen, T. Tang-Huau, K. Meade-White, B. Kaza, B.J. Smith, P. W. Hanley, J. Lovaglio, M. A. Jarvis, C. Shaia, H. Feldmann

**Affiliations:** Laboratory of Virology, Hamilton, MT, Unites States; Rocky Mountain Veterinary Branch, Division of Intramural Research, National Institute of Allergy and Infectious Diseases, National Institutes of Health; Hamilton, MT, Unites States; University of Plymouth; Plymouth, United Kingdom; The Vaccine Group Ltd; Plymouth, United Kingdom

## Abstract

The continuing emergence of SARS-CoV-2 variants calls for regular assessment to identify differences in viral replication, shedding and associated disease. In this study, African green monkeys were infected intranasally with either a contemporary D614G or the UK B.1.1.7 variant. Both variants caused mild respiratory disease with no significant differences in clinical presentation. Significantly higher levels of viral RNA and infectious virus were found in upper and lower respiratory tract samples and tissues from B.1.1.7 infected animals. Interestingly, D614G infected animals showed significantly higher levels of viral RNA and infectious virus in rectal swabs and gastrointestinal tract tissues. Our results indicate that B.1.1.7 infection in African green monkeys is associated with increased respiratory replication and shedding but no disease enhancement similar to human B.1.1.7 cases.

**One-Sentence Summary:** UK B.1.1.7 infection of African green monkeys exhibits increased respiratory replication and shedding but no disease enhancement

## Main Text

Severe acute respiratory syndrome coronavirus 2 (SARS-CoV-2) emerged in late 2019 as the causative agent of coronavirus disease 2019 (COVID-19). COVID-19 was declared a pandemic by the World Health Organization in March 2020 (*1*) and has now infected more than 170 million people with over 3.7 million deaths (*2*).

Enhanced sequence-based surveillance and epidemiological studies have led to the identification of multiple SARS-CoV-2 variants carrying distinct mutations that may impact transmissibility, disease severity and/or effectiveness of treatments and vaccines. The SARS-CoV-2 B.1.1.7 variant was first reported within the English county of Kent from the United Kingdom (UK) (*3*) and has since been classified as a ‘Variant of Concern’ (VOC) associated with increased transmissibility and potentially increased disease severity but with minimal impact on the efficacy of monoclonal antibody treatment (*4*). Several clinical reports supported the increase in transmissibility associated with the B.1.1.7 VOC with up to a 90% increase in transmission compared to earlier variants (*5-7*). However, the reported increase in mortality of the B.1.1.7 VOC (*8-10*) seen from earlier COVID case analysis has recently been questioned (*11-12*). Apart from clinical studies, experimental infections in animals - ideally in species closely related to humans such as nonhuman primates (NHPs) - is one way to assess transmissibility and disease severity of emerging SARS-CoV-2 variants.

The rhesus macaque model of SARS-CoV-2 infection was established early in the pandemic (*13-15*) and has been used to test SARS-CoV-2 therapeutics and vaccines (*16-19*). Additional NHP species, such as cynomolgus macaques, baboons and marmosets have been investigated for their susceptibility to SARS-CoV-2 in an attempt to develop models exhibiting increased disease severity (*20, 21*). None of these models result in severe disease, but susceptible NHP species exhibit oral and nasal shedding and develop mild to moderate respiratory disease in the upper and lower respiratory tract. An African green monkey (AGM) model of SARS-CoV-2 infection has recently been developed, wherein animals show greater severity of disease (*22, 23*), with intranasal infection of AGMs resulting in significant shedding and respiratory disease (*24*). The AGM model may thereby represent a more natural NHP model for SARS-CoV-2 infection and disease.

In our current study, the AGM intranasal model of SARS-CoV-2 infection was used to assess differences between a contemporary SARS-CoV-2 D614G variant, which was circulating in the summer of 2020, and the B.1.1.7 VOC that emerged in the UK in late 2020. Herein, we report and discuss differences in organ tropism, replication kinetics and shedding between the two SARS-CoV-2 variants.

### Infection with B.1.1.7 was not associated with a significant increase in disease severity in the AGM model

Following intranasal infection with 1×10^6^ infectious particles of either the SARS-CoV-2 D614G (n=5) or the B.1.1.7 variant (n=6) (5×10^5^ per naris) using a nasal atomization device, animals were monitored and scored daily for clinical signs of disease including changes in general appearance, respiration, food intake and fecal output and locomotion. Clinical signs were mild with both groups of AGMs displaying only minor changes in respiration and showing reduced appetite that negatively impacted volume of feces produced. Slight differences were observed between the two variant groups. B.1.1.7 infected animals had elevated scores early in infection peaking at 2 days post-infection (dpi), which subsequently returned toward baseline (Fig 1A). In contrast, scores for the D614G animals increased slowly peaking at 4dpi and remained stable until euthanasia (Fig 1A, Table S1). Radiographs were taken at each examination and scored for pulmonary infiltrates, progression of which were similar to the clinical scores (Fig 1B, Table S2). B.1.1.7 infected AGMs scored slightly higher earlier and peaked at 1dpi, whereas D614G infected animals scored higher later and peaked at 3dpi. Overall, the changes in clinical and radiographic scores were not significantly different between the two groups even though minor differences were noted in disease progression.

**Figure 1:**
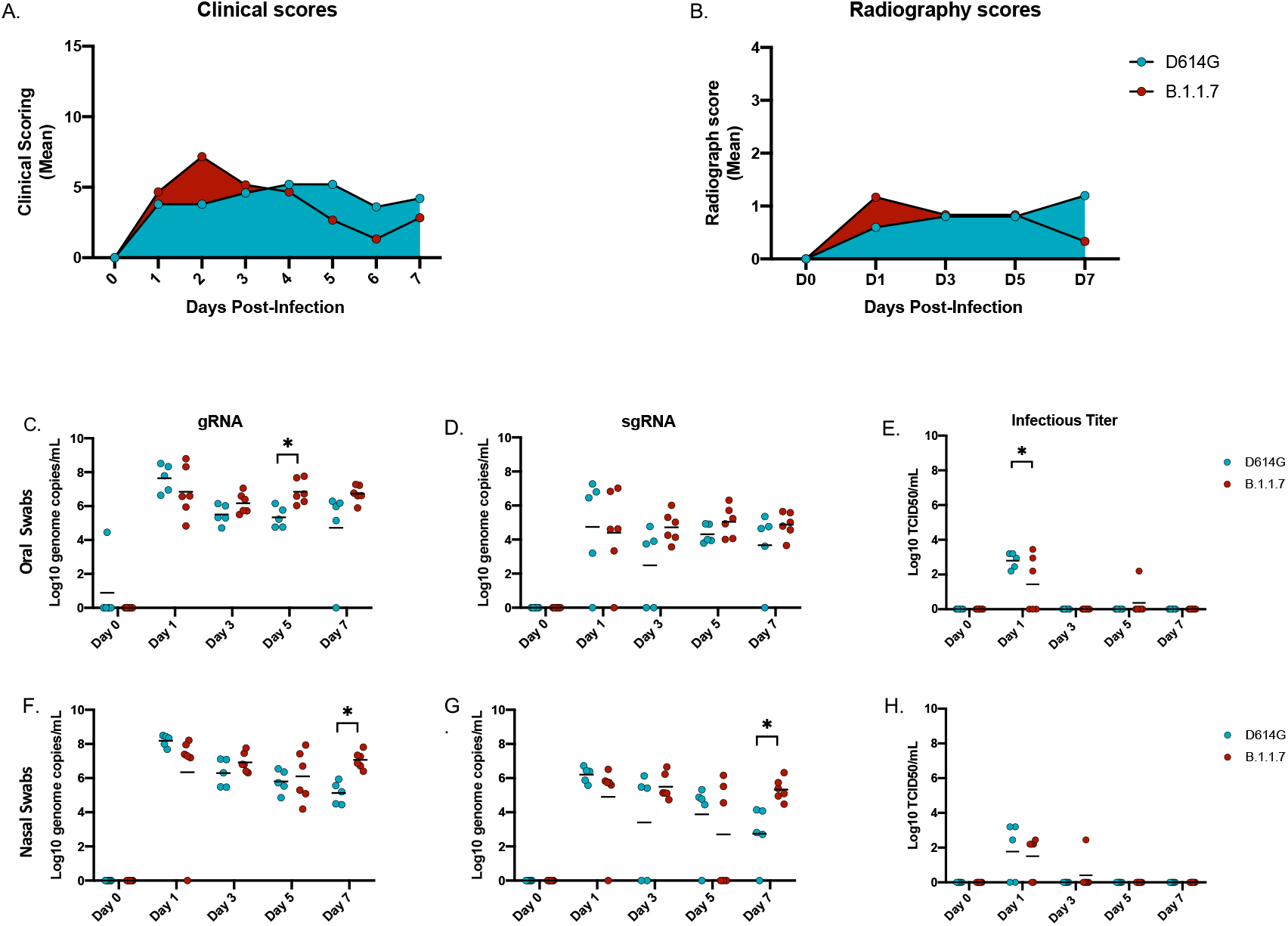
Clinical scoring, radiographs and oral and nasal shedding. AGMs were infected with either the D614G or B.1.1.7 SARS-CoV-2 variant intranasally utilizing the Nasal Mucosal Atomization Device. (A) AGMs were scored daily for clinical signs of disease including changes in general appearance, respiration, food intake and feces as well as locomotion (B) Radiographs were taken on clinical exam days (0, 1, 3, 5, 7) and scored for pulmonary infiltrates. Swabs were taken on clinical exam days (0, 1, 3, 5 and 7) and used as a correlate for virus shedding. Viral RNA total genome (gRNA) and subgenome (sgRNA) copies were determined by qRT-PCR. Infectious virus was titered on VeroE6 cells. (C-E) Viral shedding in oral swabs. Statistical significance was found at day 5 in total RNA (C, p-value <0.05) and at day 1 in infectious titers (E, p-value <0.05). (F-H) Viral shedding in nasal swabs. Statistical significance was found at day 7 in gRNA (F, p-value <0.05) and at day 7 sgRNA (G, p-value <0.05). Multiple t-tests were used to compare the gRNA, sgRNA and infectious titers between groups.

### B.1.1.7 replication/shedding from the upper respiratory tract was increased compared to D614G

Oral and nasal swabs were taken at each examination to assess virus replication in the upper respiratory tract and virus shedding. SARS-CoV-2 RNA was measured with qPCR assays targeting either total viral RNA (gRNA, N assay) or subgenomic viral RNA (sgRNA, E assay) (Fig 1). Total gRNA in oral swabs was significantly higher at 5dpi in B.1.1.7 compared to D614G infected animals; this difference between variants was maintained but dropped below significance by 7dpi (Fig 1C). There were no significant differences in sgRNA levels in oral swabs at any time point, but levels consistently trended higher in the B.1.1.7 infection group starting at 3dpi (Fig 1D). Although viral RNA was detectable throughout the study, infectious virus was only isolated from oral swabs at 1dpi with significantly higher titers for the D614G infected animals (Fig 1E). Consistent with the oral swabs, higher levels of viral RNA were detected in nasal swabs collected from AGMs infected with B.1.1.7. By 7dpi these animals were all shedding significantly more viral RNA than those infected with D614G (Fig 1F, G). Infectious virus in the nasal swabs was recovered predominantly at 1dpi and there was no difference between groups (Fig 1H).

### B.1.1.7 replication in the lower respiratory tract was increased compared to D614G

Samples to assess viral replication kinetics in the lower respiratory tract were collected with broncho cytology brushes (BCB) at 3, 5 and 7dpi and bronchoalveolar lavage (BAL) at 3 and 5dpi. Total viral RNA levels were consistent between both sampling methods (∼10^5^-10^7^ genome copies/ml) at each day sampled (Fig 2A-C; Fig S1). The gRNA levels were consistently higher in AGMs infected with B.1.1.7 and significantly increased at the final time points of BCB (7dpi) (Fig 2A) and BAL (5dpi) sampling (Fig S1). Although not significantly different at any time point, sgRNA was higher in the B.1.1.7 AGMs at 7dpi in the BCB and in BAL at the final day of collection (5dpi) (Fig 1B; Fig S1). Higher levels of infectious virus were also isolated from animals infected with B.1.1.7, particularly in BCB samples (Fig 2C; Fig. S1).

**Figure 2:**
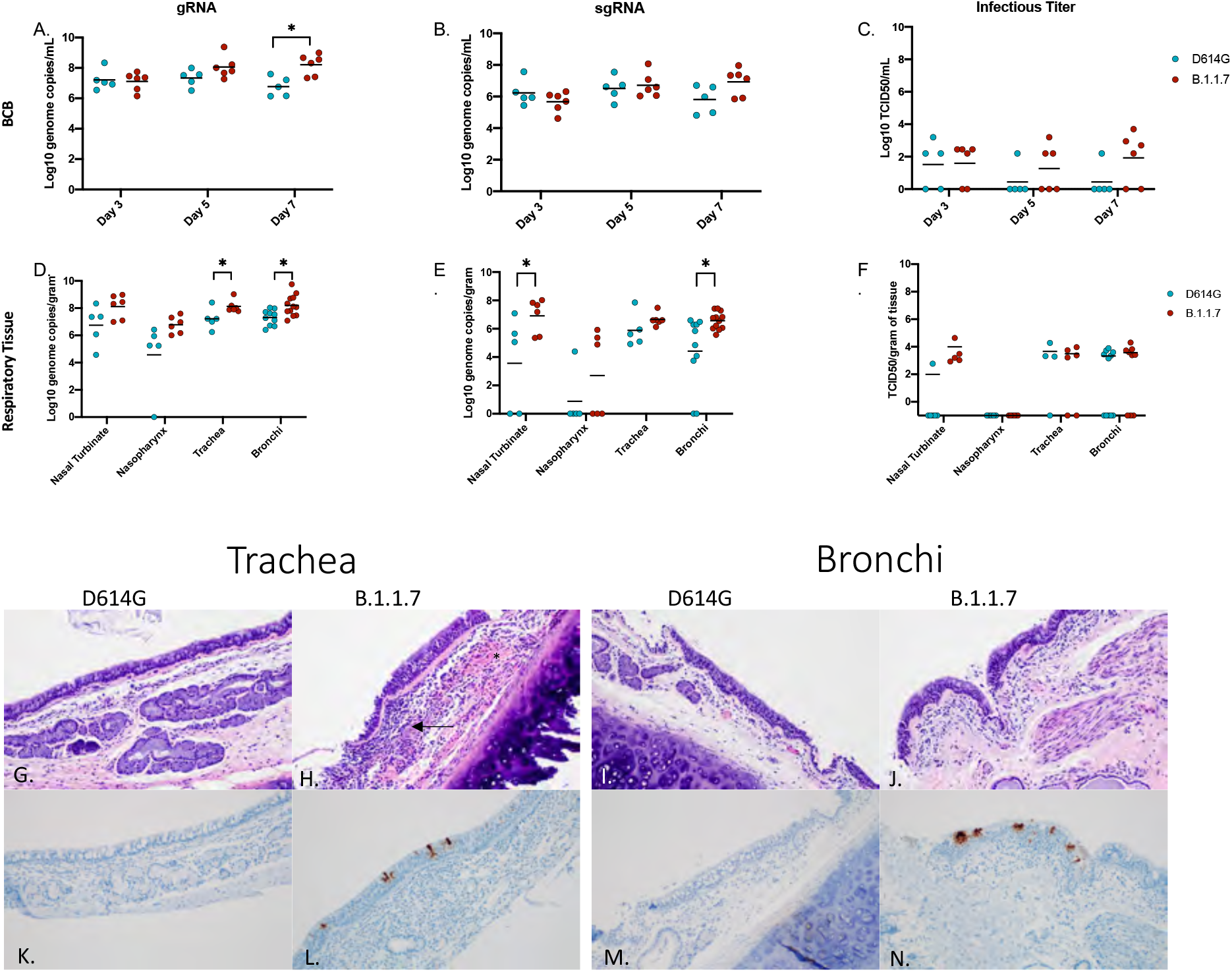
Virus load in the lower respiratory tract. AGMs were infected with either the D614G or B.1.1.7 SARS-CoV-2 variant intranasally utilizing the Nasal Mucosal Atomization Device. Bronchial cytology brush (BCB) samples were collected on days 3, 5 and 7 post-infection and samples were analyzed for gRNA, sgRNA and infectious virus. (A-C) A significant difference in gRNA collected in the BCBs was detected 7 days-post-infection (A, p-value <0.05). Animals were euthanized on day 7 post-infection and respiratory tissues were collected for analyses for gRNA, sgRNA and infectious virus (D-F). gRNA was significantly different in the trachea and bronchi (D, p-value <0.05). sgRNA differed significantly in the nasal turbinate and bronchi (E, p-values <0.05). Infectious virus did not differ significantly in any tissue (C). Multiple t-tests were used to compare tissue between groups. Pathology and immunoreactivity in the trachea and bronchi (G-N). (G, H) Normal trachea found in the D614G vs. trachea with cellular infiltrates, hemorrhage (*) and fibrin (arrow) found in the submucosa in the B.1.1.7. (K, L) immunoreactivity in the trachea of D614G vs B.1.1.7. (I, J) Normal bronchi found in the D614G vs bronchi with inflammation and cellular infiltrates found in the B.1.1.7. (M, N) Immunoreactivity in bronchi of the D614G vs B.1.1.7. (HE G, H; IHC K, L 100x; HE I, J; IHC M, N 100x).

Post-mortem tissues were collected at 7dpi for virological analysis and pathology. Respiratory tissues including nasal turbinate, nasopharynx, trachea and left and right bronchi and a section from each lung lobe were examined for viral RNA and infectious virus. Total and sgRNA was higher in all respiratory tissues in the B.1.1.7 infected animals and was significantly higher in the trachea and bronchi (Fig 2D,E). Although not statistically significant, AGMs infected with B.1.1.7 had higher levels of infectious virus in nasal turbinates, trachea and bronchi (Fig 2F). Consistent with the viral RNA and infectious titer results, most animals (5 out of 6) infected with B.1.1.7 had inflammation of the trachea, with only 1 of 5 animals infected with the D614G variant having any similar observable lesion. Similarly, lesions were found in 8 of 10 bronchi from B.1.1.7 infected AGMs compared to only 2 of 8 of D614G animals, with lesions corresponding to SARS-CoV-2 immunoreactivity by immunohistochemistry (IHC) (Fig 2G-N).

Total and sgRNA was significantly higher in the lungs of B.1.1.7-compared to D614G-infected animals (Fig 3A, B); however, elevated levels of infectious virus in B.1.1.7 animals remained just below statistical significance (Fig 3C). Lung lesions for both groups were minor but consistent with SARS-CoV-2 pneumonia and included thickening and inflammation of alveolar septa and the presence of fibrin (Fig 3D-G). SARS-CoV-2 immunoreactivity by IHC was also limited and not directly associated with foci of inflammation (Fig 3H-K).

**Figure 3:**
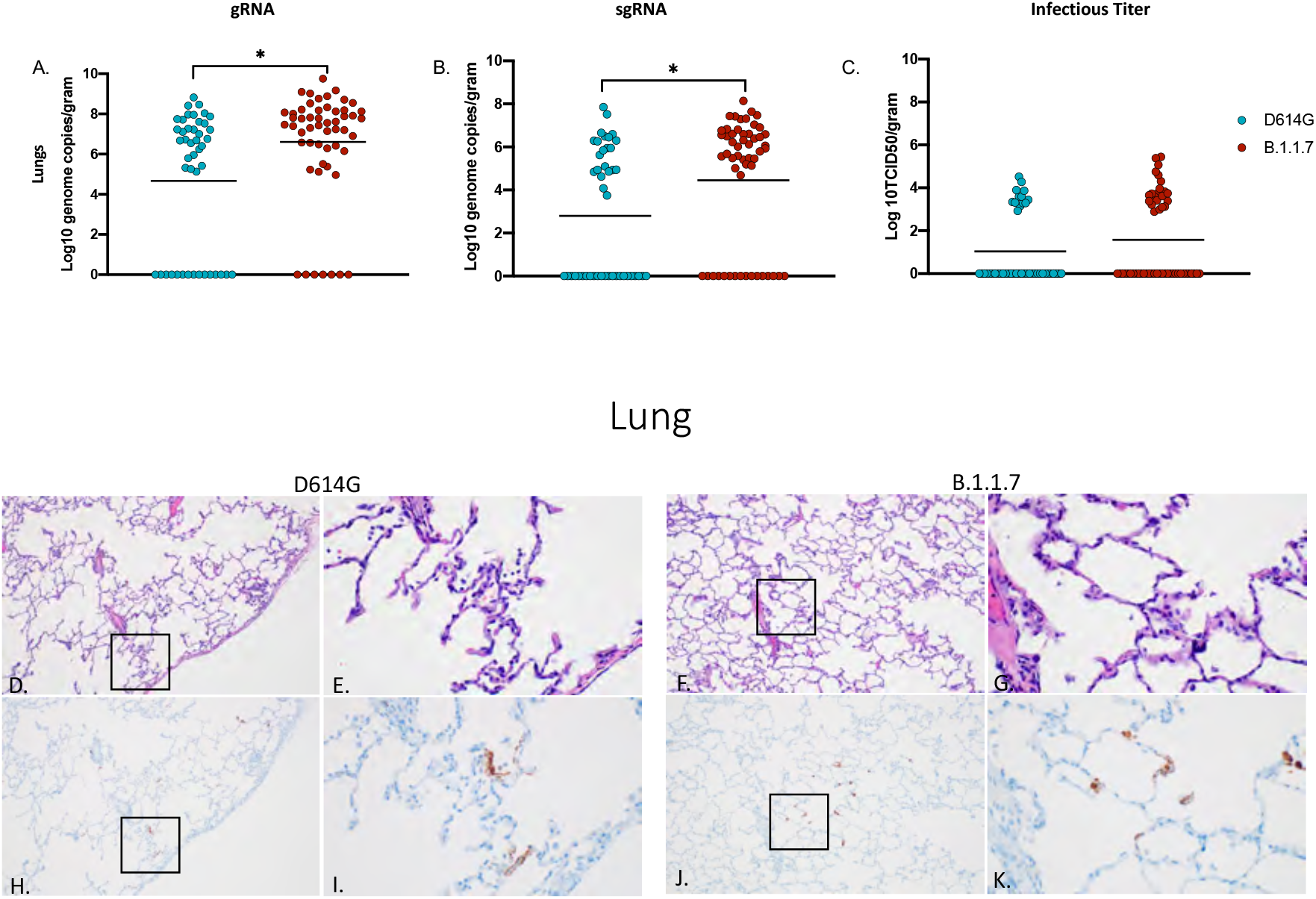
Virus load, pathology and immunoreactivity in the lungs. Animals were euthanized on day 7 post-infection and a section from each lung lobe were collected and analyzed for gRNA, sgRNA and infectious virus. Results of each assay were combined to look at the lungs in total (A-C). Significant differences were detected in lungs in both gRNA and sgRNA (D, p-value <0.05 and E. p-value <0.05) but not in infectious virus (C). Multiple t-tests were used to compare the two groups for statistical significance. Pathology and immunoreactivity in the lungs (D-K). (D, E) Minimally thickened and inflamed alveolar septa with multifocal pneumocyte immunoreactivity in the D614G and B.1.1.7 samples (HE D, F; IHC H, J 100x; HE E, G; IHC I, K 400x).

### D614G replication in the gastrointestinal (GI) tract was increased compared to B.1.1.7

Total viral gRNA was detected in cervical lymph nodes, tonsil, heart, liver, spleen, ileum and cecum in both groups (Fig 4A). With the exception of one liver sample, sgRNA was not detected in liver, spleen nor kidneys (Fig 4B). Notably and in marked contrast to respiratory tissues, AGMs infected with D614G had significantly more total and sgRNA in the ileum and cecum than B.1.1.7 infected AGMs (Fig 4A, B). Levels of viral RNA corresponded to infectious virus with only D614G animals having detectable infectious SARS-CoV-2 in these two GI-derived tissues (Fig 4C). Similarly, gRNA in rectal swabs peaked and was significantly higher at 7dpi in D614G infected animals (Fig 4D). sgRNA was recovered intermittently across the study (Fig 4E), but infectious virus was recovered from rectal swabs of only one D614G animal (Fig 4F). Viral replication in the ileum was associated with inflammation in D614G infected AGMs and corresponded with detectable viral antigen by IHC (Fig 4G,H,K,L), while AGMs infected with the B.1.1.7 had no observable inflammation or viral antigen (Fig 4I,J,L,M). In D614G infected animals, only one AGM presented with inflammation and a small amount of associated viral antigen in the cecum. No infectious virus was isolated from any of the other non-respiratory tissue in either group and pathology was unremarkable in these tissues.

**Figure 4:**
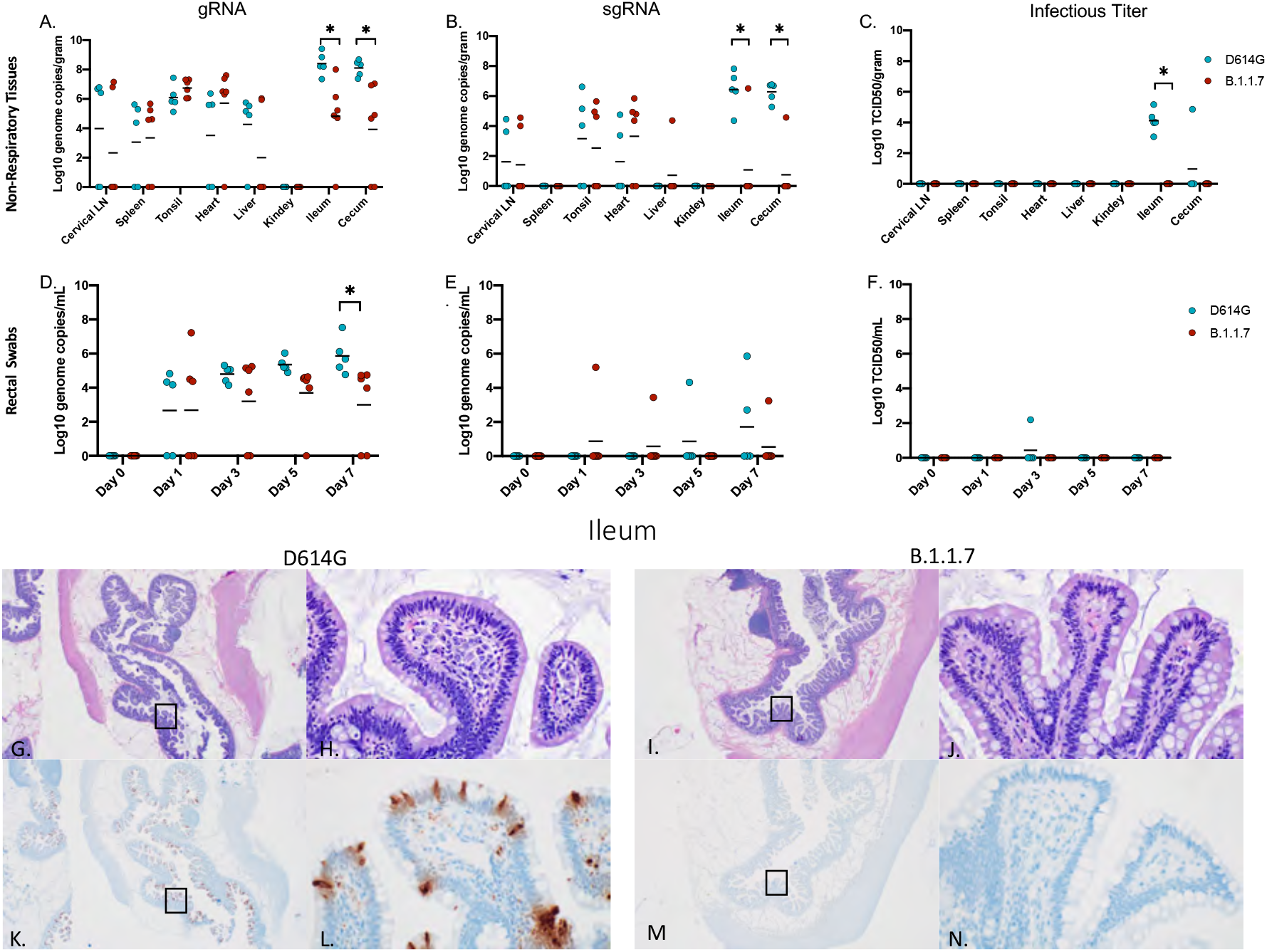
Viral load in gastrointestinal tract, pathology and immunoreactivity. AGMs were euthanized 7 days post-infection and tissues were collected to determine viral load, pathology and immunoreactivity (A-C). gRNA and sgRNA was significantly different in the ileum and cecum (A, p-value <0.05 and B, p-value <0.05). Infectious virus was significantly different in the ileum (C, p-value <0.05), no other tissues were significantly different. (D-F) Viral shedding in rectal swabs. Statistical significance was found at day 7 in total RNA (D, p-value <0.05). tatistical significances was determined by multiple t-tests between the two groups. Pathology and immunoreactivity in the ileum (G-N). (G, H) Normal mucosa with multifocal mucosal immunoreactivity in the D614G challenged ileum (HE G, IHC K, 20x; HE H, IHC L, 400x). (I, J) Normal mucosa and no immunoreactivity in the B.1.1.7 challenged ileum (HE I, IHC M, 20x; HE J, IHC N, 400x).

### Infection with neither variant was associated with marked changes in hematology, blood chemistry and coagulation, nor a broad systemic cytokine response

Blood and serum samples were collected for hematology, blood chemistry, coagulation assays and cytokine analysis at every clinical examination. No differences were found in the hematology (Fig S2A-L), blood chemistry (Fig S2M-T) or coagulation assays (Fig S3) between the D614G and B.1.1.7 infected AGMs. Of the cytokines examined (Fig. S4), IL6 was the only pro-inflammatory cytokine that was significantly different between the groups, with IL6 being elevated in the D614G group at 3dpi and 5dpi compared to B.1.1.7 infected animals (Fig S4A). Levels of T cell chemo-attractants IP-10 (CXCL 10) (Fig S4B) and I-Tac (CXCL 11) (Fig S4) were also increased at 1dpi in the D614G group but were not sustained throughout the study.

The regular emergence of SARS-CoV-2 variants represents a constant public health challenge with the COVID-19 pandemic. Although epidemiological and clinical data can give insight into characteristics of a new variant, this data can be limited initially and may be biased by many factors. Animal models provide the ability to directly compare biological and clinical characteristics of multiple SARS-CoV2 variants in a study with limited variables providing invaluable data otherwise unattainable.

In the present study, we have used the AGM intranasal infection model to compare the B.1.1.7 VOC, a variant that emerged in the UK in September of 2020 and then quickly spread throughout the world (*5, 7*), with a contemporary D614G variant, in terms of virus replication, shedding and disease severity. A nasal atomization device was used to infect NHPs to most closely mimic natural infection in human. Following infection with either variant, animals in both groups exhibited minor differences in disease progression but overall disease signs were similar with mild respiratory disease for both B.1.1.7 and D614G (Fig. 1A,B; Table S1 & S2). Increased disease severity was initially reported for human B.1.1.7 cases (*8-10*), but more recent studies have contradicted these earlier claims (*11, 12*). The outcome of our study using a NHP surrogate model supports findings from these more recent studies indicating that B.1.1.7 VOC is not associated with increased disease severity.

Although no animals in this study developed severe disease, our analysis reveal differences between the two variants in terms of their replication within the respiratory system. Viral RNA and infectious virus in the lower respiratory tract tissues were more prevalent in the animals infected with B.1.1.7 compared to D614G, especially at later timepoints suggesting the development of a stronger respiratory component associated with the emerging VOC (Figs 2, Fig 3). Consistent with higher levels of viral replication in the upper respiratory tract, shedding of viral RNA in both the nose and oral cavity was also higher in B.1.1.7 infected animals (Fig. 1), which support reports from human infection data showing that the B.1.1.7 VOC is more transmissible than earlier variants (*6, 7, 25*).

The pathology associated with infections differed between the two variants. B.1.1.7 replicated at higher levels in the respiratory tract resulting in lesions that were both more numerous and severe than seen for D614G infected animals. In contrast, D614G replicated at higher levels in the GI tract and the associated pathology seen in these animals correlated with this difference in GI replication. This finding was supported by higher levels of viral RNA in rectal swabs indicating the possibility of fecal transmission (Fig. 4). This may indicate that D614G is more suited to replication in the digestive tract than other variants which is in line with clinical studies conducted in early to mid-2020 that reported GI symptoms in approximately 15-20% of COVID-19 patients (*26*). The difference observed in these studies may also indicate that genetic alterations in B.1.1.7 may have not merely resulted in a general increased rate of replication but may have also altered organ tropism. Clinical studies concerning B.1.1.7 have to date been mainly focused on respiratory symptoms (*8-11*). However, with clinical studies still underway it remains to be seen whether the observed changes in tissue tropism and replication detected here in the AGM model will correspond to a drop in reported GI symptoms in those infected with the B.1.1.7 VOC compared to infections with contemporary SARS-CoV-2. It is also possible that AGMs may be more prone to GI tract infections and an earlier SARS-CoV-2 study with AGMS using the nCoV-WA1-2020 isolate suggests this could be true. That study had a single animal that exhibited infection in the GI tract up to 10dpi but the remaining animals did not (*27*). Future studies should address potential differences in organ tropism associated with SARS-CoV-2 variants.

A recent study posted on a preprint server examining SARS-CoV-2 variants in the rhesus macaque model showed no difference in viral replication nor disease between the D614G and B.1.1.7 variants (*28*). Notably, this study used double the infectious dose via intranasal challenge using the same atomizing device, but also added an intratracheal inoculation route (*28*). Although AGMs and rhesus macaques share similar ACE-2/RBD binding affinities (*29*), we cannot rule out the different NHP species as a potential factor in the outcome of infection. The marginally higher dose and additional route of infection may have been additional factors affecting infection outcomes.

In conclusion, our results from the intranasal AGM COVID-19 surrogate model support the most recent data from B.1.1.7 in humans, providing direct empirical data for increased replication in respiratory tissue, but with no enhancement of disease. Although the lack of clear statistical significance for some of the parameters may be regarded as possible limitation of the study, the use of these multiple parameters to independently verify one another addresses this concern. One way to obtain statistical significance more uniformly is to increase NHP numbers something that is ethically sensitive and difficult given the current issues with NHP availability. NHPs remain a surrogate model for humans and results presented herein provide direct experimental evidence that support recent clinical observations, which indicate that the B.1.1.7 VOC has characteristics of increased replication in respiratory tissues with enhanced shedding from the nose and oral cavity resulting in advanced transmission. A further notable and interesting observation of differences between these two variants in terms of distinct viral tissue/organ tropism warrants further attention as such changes would have public health implications in terms of transmission and disease manifestation.

## Acknowledgements

The authors are thankful to the animal caretakers and histopathology group of the Rocky Mountain Veterinary Branch (NIAID, NIH) for their support with animal related work, to the Research Technologies Branch (NIAID, NIH) for sequencing of stock viruses, and Anita Mora (NIAID, NIH) for help with the display items. We are grateful to Emmie de Wit and Vincent Munster for their discussions and help with virus stock preparations. The B.1.1.7 variant was obtained through BEI Resources (Bassam Hallis, Sujatha Rashid), NIAID (Ranjan Mukul, Kimberly Stemple) and the NIH.

## Author contributions

Conceptualization: KR, MAJ, HF; Methodology: KR, FF, AO, FH, TTH, KMW, BK, BJS, PWH, JL, CS; Visualization: KR, FH; Funding acquisition: HF; Project administration: KR, HF; Supervision: KR, PWH, CS, HF; Writing (original draft): KR, MAJ, CS, HF; Writing (review & editing): FF, AO, FH, TTH, KMW, BK, BS, PWH, JL, CS

## Data and materials availability

All data are available in the main text or the supplementary materials. Additional information can be requested through the corresponding author.

## Disclaimer

The opinions, conclusions and recommendations in this report are those of the authors and do not necessarily represent the official positions of the National Institute of Allergy and Infectious Diseases (NIAID) at the National Institutes of Health (NIH). There was no conflict in interest identified for any individual involved in the study.

## Supplementary Materials

## Materials and Methods

### Biosafety and ethics

All SARS-CoV-2 studies were approved by the Institutional Biosafety Committee (IBC) and performed in high biocontainment (BSL3/BSL4) at Rocky Mountain Laboratories (RML), NIAID, NIH. All sample processing in high biocontainment and sample removal followed IBC-approved Standard Operating Protocols (SOPs) (*1*). All experiments involving AGMs were performed in strict accordance with approved Institutional Animal Care and Use Committee protocols and following recommendations from the Guide for the Care and Use of Laboratory Animals of the Office of Animal Welfare, National Institutes of Health and the Animal Welfare Act of the US Department of Agriculture, in an Association for Assessment and Accreditation of Laboratory Animal Care International (AAALAC)-accredited facility. AGMs were placed in a climate-controlled room with a fixed 12-hour light-dark cycle. Animals were singly housed in adjacent primate cages allowing social interactions and provided with commercial monkey chow, treats, and fruit twice daily with water *ad libitum*. Environmental enrichment was provided with a variety of human interaction, manipulanda, commercial toys, movies, and music. AGMs were monitored at least twice daily throughout the study.

### Virus and cells

SARS-CoV-2 isolate SARS-CoV-2/human/USA/RML-7/2020 (MW127503.1), strain D614G, was obtained from a nasopharyngeal swab obtained on July 19, 2020. Sequencing of the viral stock showed it to be 100% identical to the deposited Genbank sequence and no contaminants were detected (*2*). SARS-CoV-2 variant B.1.1.7 (hCoV-19/England/204820464/2020, EPI_ISL_683466) was obtained from Public Health England via BEI Resources (Manassas, VA, USA). The supplied passage 2 material was propagated once in Vero E6 cells. Sequencing confirmed the presence of three SNPs in this stock: nsp6 D156G (present in 14% of all reads), nsp6 L257F (18%) and nsp7 V11I (13%) (*3*).

Virus propagation was performed in Vero E6 cells in DMEM (Sigma-Aldrich, St Louis, MO, USA) supplemented with 2% fetal bovine serum, 1 mM L-glutamine, 50 U/ml penicillin and 50 μg/ml streptomycin (DMEM2). Vero E6 cells were maintained in DMEM supplemented with 10% fetal bovine serum, 1 mM L-glutamine, 50 U/ml penicillin and 50 μg/ml streptomycin (DMEM10). Mycoplasma testing of cell lines and viral stocks is performed regularly, and no mycoplasma was detected.

### Study design

Eleven SARS-CoV-2 seronegative AGMs (3.8-6.7 kg) were divided into 2 groups for infection with either the contemporary D614G strain (RML7) (n=5) or the recently emerged (UK variant) (n=6). A Nasal Mucosal Atomization Device (Teleflex, MAD110) was used to deliver 10^6^ infectious particles (5×10^5^ per naris diluted in 500ul DMEM with no additives). Clinical examinations were performed on days 0, 1, 3, 5 and 7. Blood and serum were collected for hematology, blood chemistry, coagulation and virological analysis. Oral, nasal and rectal swabs were collected at every examination for virological analysis. Bronchial cytology brushes were collected on days 3, 5 and 7 and bronchioalveolar lavage (BAL) samples were also collected on days 3 and 5 for virological analysis. Tissues were collected following euthanasia on day 7 for pathology and virological analysis. Studies were performed in successive weeks and different animal study groups to avoid contamination between studies, the D416G study was run first followed by the B.1.1.7 study.

### Virus titration

Virus isolation was performed on tissues following homogenization in 1 mL DMEM using a TissueLyser (Qiagen, Germantown, MD, USA) and inoculating Vero E6 cells in a 96 well plate with 200 µL of 1:10 serial dilutions of the homogenate. One hour following inoculation of cells, the inoculum was removed and replaced with 200 µL DMEM. Virus isolation of blood and swab samples were performed in a similar manner. Samples were vortexed for 30 seconds before performing the 1:10 dilution series. The inoculum (200ul) was placed on cells and rocked for 1h. Infectious supernatant was removed and replaced with fresh DMEM. Seven days following inoculation, cytopathogenic effect was scored and the TCID50 was calculated using the Reed-Muench formula (*4*).

### Viral RNA detection

qPCR was performed on RNA samples extracted from swabs or tissues using QiaAmp Viral RNA or RNeasy kits, respectively (Qiagen, Germantown, MD, USA). Viral RNA was detected with one-step real-time RT-PCR assays designed to amplify total viral RNA (N gene) (*5*) or sgRNA by amplifying a region of E gene to detect replicating virus (*6*). Dilutions of RNA standards counted by droplet digital PCR were run in parallel and used to calculate viral RNA genome copies. A Rotor-Gene probe kit (Qiagen, Germantown, MD, USA) was used to run the PCRs according to the instructions of the manufacturer.

### Hematology, Serum Chemistry and Coagulation

Hematology analysis was completed on a ProCyte DX (IDEXX Laboratories, Westbrook, ME, USA) and the following parameters were evaluated: red blood cells (RBC), hemoglobin (Hb), hematocrit (HCT), mean corpuscular volume (MCV), mean corpuscular hemoglobin (MCH), mean corpuscular hemoglobin concentration (MCHC), red cell distribution weight (RDW), platelets, mean platelet volume (MPV), white blood cells (WBC), neutrophil count (abs and %), lymphocyte count (abs and %), monocyte count (abs and %), eosinophil count (abs and %), and basophil count (abs and %). Serum chemistries were completed on a VetScan VS2 Chemistry Analyzer (Abaxis, Union City, CA, USA) and the following parameters were evaluated: glucose, blood urea nitrogen (BUN), creatinine, calcium, albumin, total protein, alanine aminotransferase (ALT), aspartate aminotransferase (AST), alkaline phosphatase (ALP), total bilirubin, globulin, sodium, potassium, chloride, and total carbon dioxide. Coagulation values were determined from citrated plasma utilizing a STart4 Hemostatis Analyzer and associated testing kits (Diagnostica Stago, Parsippany, NJ, USA).

### Cytokine analyses

Concentrations of cytokines and chemokines present in the serum from SARS-CoV-2 infected AGMs were quantified using a multiplex bead-based assay (1:4 dilution)-the LEGENDPlex Non-Human Primate Cytokine/Chemokines 13-plex (BioLegend, San Diego, CA USA). Analytes detected by this panel are the following: IFN-γ, IL-1β, IL-6, IL-8, MCP-1, MIP-1α, MIP-1β, MIG, TNF-α, I-TAC, RANTES, IP-10, and Eotaxin. Samples were diluted 1:4 in duplicate prior to processing according the manufacturer’s instructions. Samples were read using the BD FACS Symphony instrument (BD Biosciences, San Jose, CA USA) and analyzed using LEGENDplexTM Data Analysis Software following data acquisition.

### Thoracic radiographs

Ventro-dorsal and right/left lateral radiographs were taken on clinical exam days prior to any other procedures (e.g. bronchoalveolar lavage, nasal flush). Radiographs were evaluated and scored for the presence of pulmonary infiltrates by two board-certified clinical veterinarians according to a previously published standard scoring system (*7*). Briefly, each lung lobe (upper left, middle left, lower left, upper right, middle right, lower right) was scored individually based on the following criteria: 0 = normal examination; 1 = mild interstitial pulmonary infiltrates; 2 = moderate interstitial pulmonary infiltrates, perhaps with partial cardiac border effacement and small areas of pulmonary consolidation (alveolar patterns and air bronchograms); and 3 = pulmonary consolidation as the primary lung pathology, seen as a progression from grade 2 lung pathology. At study completion, thoracic radiograph findings were reported as a single radiograph score for each animal on each exam day. To obtain this score, the scores assigned to each of the six lung lobes were added together and recorded as the radiograph score for each animal on each exam day. Scores range from 0 to 18 for each animal on each exam day.

### Histology and Immunohistochemistry

Tissues were fixed in 10 % neutral buffered formalin with two changes, for a minimum of 7 days according to an IBC-approved SOP. Tissues were processed with a Sakura VIP-6 Tissue Tek, on a 12-hour automated schedule, using a graded series of ethanol, xylene, and PureAffin. Embedded tissues were sectioned at 5 μm and dried overnight at 42°C prior to staining with hematoxylin and eosin. Specific staining was detected using SARS-CoV/SARS-CoV-2 nucleocapsid antibody (Sino Biological cat#40143-MM05) at a 1:1000 dilution. The tissues were processed for immunohistochemistry using the Discovery Ultra automated stainer (Ventana Medical Systems) with a ChromoMap DAB kit (Roche Tissue Diagnostics cat#760–159) (Roche Diagnostics Corp., Indianapolis, IN, USA).

### Statistical analyses

Statistical analysis was performed in Prism 8 (GraphPad, San Diego, CA, USA). Multiple t-tests were used to assess statistical significance between the two infection groups.

## Supplementary Figures

**Figure S1:**
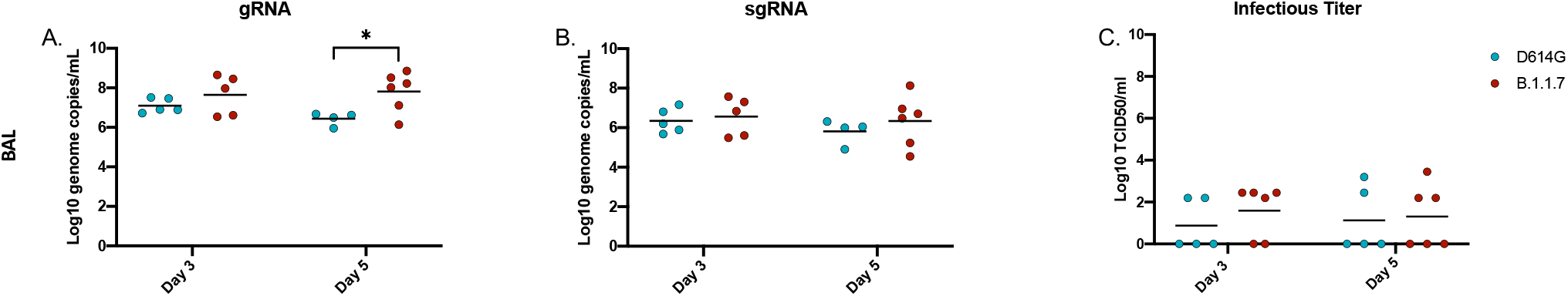
Viral loads in the lower respiratory system (BAL). AGMs were infected with either the D614G or B.1.1.7 SARS-CoV-2 variant intranasally utilizing a Nasal Mucosal Atomization Device. Bronchioalveolar lavage (BAL) samples were collected on days 3 and 5 post-infection and measured for gRNA, sgRNA and infectious titers. (A-C) A significant difference in gRNA collected in BAL samples was detected on day 5 post-infection (*p-value <0.05), no other significant differences were detected. Multiple t-tests were used to compare groups.

**Figure S2:**
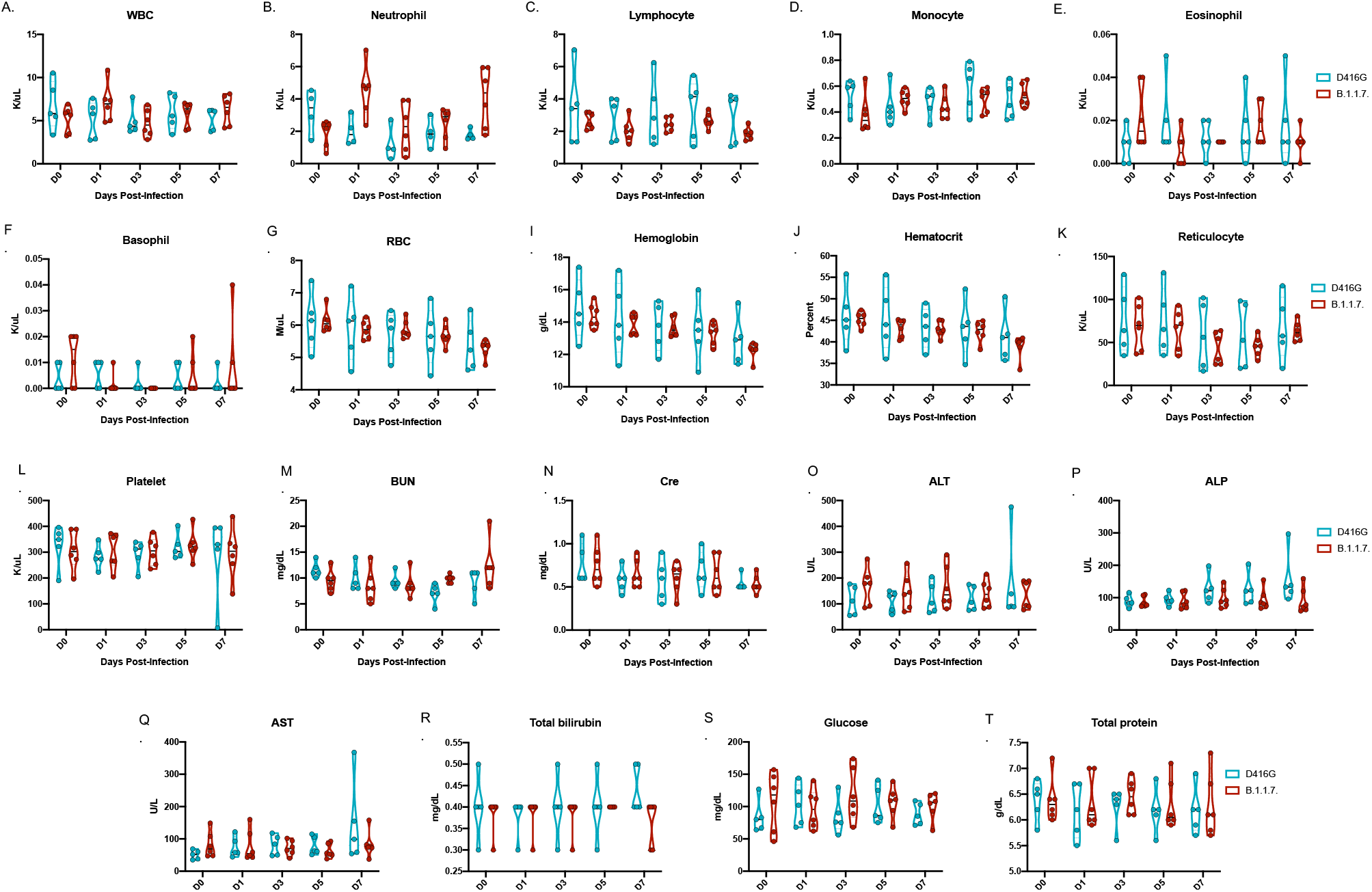
Hematology and blood chemistry following infection. Whole blood and serum samples were collected at each exam time point (days 0, 1, 3, 5 and 7) for hematology (A-L) and blood chemistry analyses (M-T). No significant changes were found in hematology (A-L), nor in blood chemistry (M-T).

**Figure S3:**
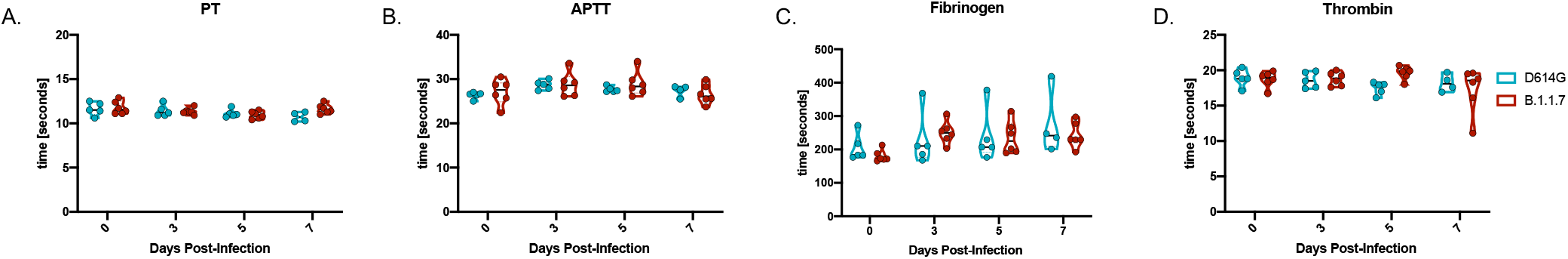
Coagulation assays following infection. Plasma samples were collected at each clinical time point (days 0, 1, 3, 5 and 7) to evaluate coagulation parameters between infected animals (A-D). No significant changes were found in PT (A), APTT (B), fibrinogen (C) or thrombin (D).

**Figure S4:**
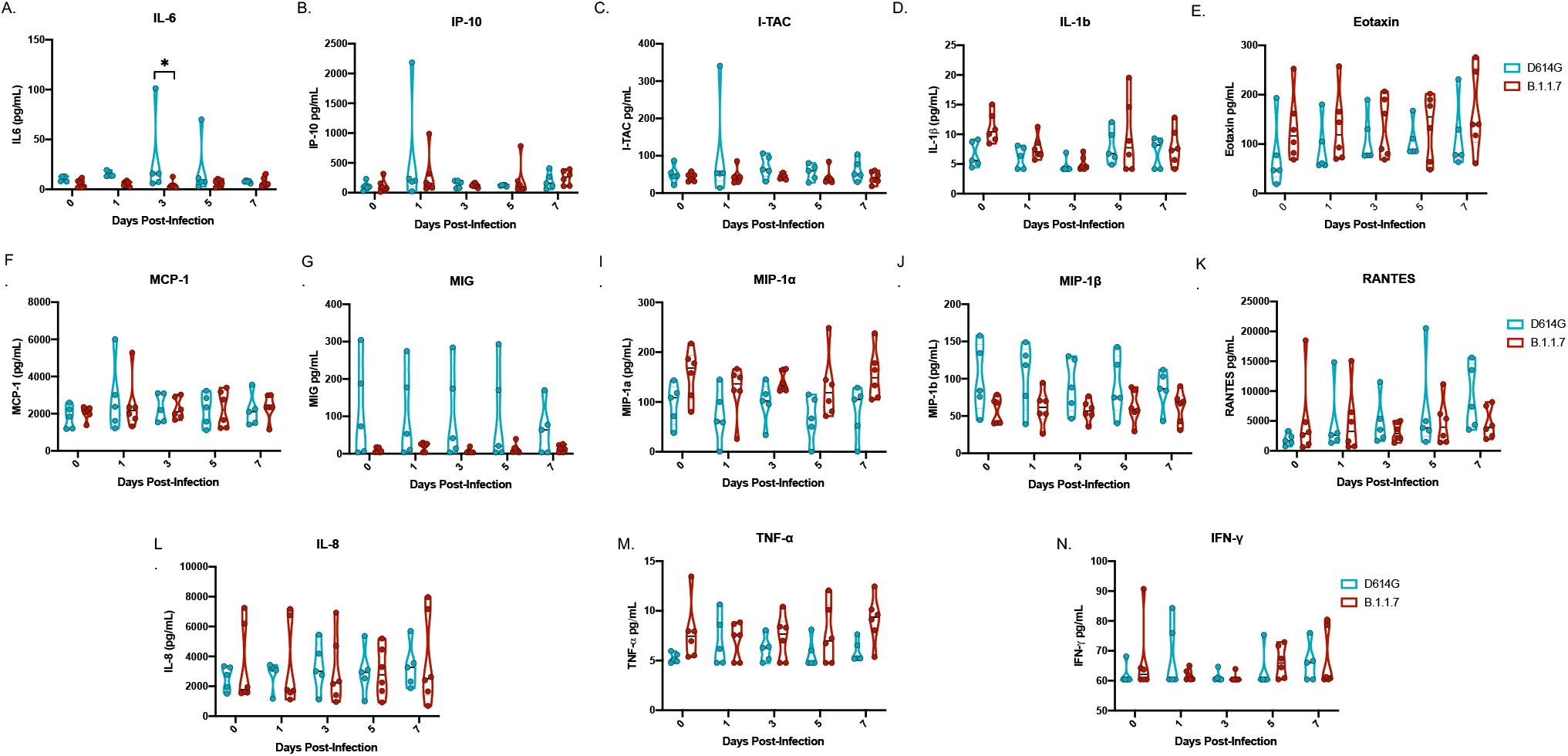
Cytokine analyses following infection. Serum was collected on days 0, 1, 3, 5 post-infection for cytokine analyses. Three notable changes were detected. Levels of IL-6 were significantly different 3 days post-infection between the two groups (*p-value 0.05) (A). Differences at 1 day post-infection were noted in both IP-10 (B) and I-TAC (C) but were not significant. Samples were analyzed by 2-way ANOVA to determine significance.

**Table S1:**
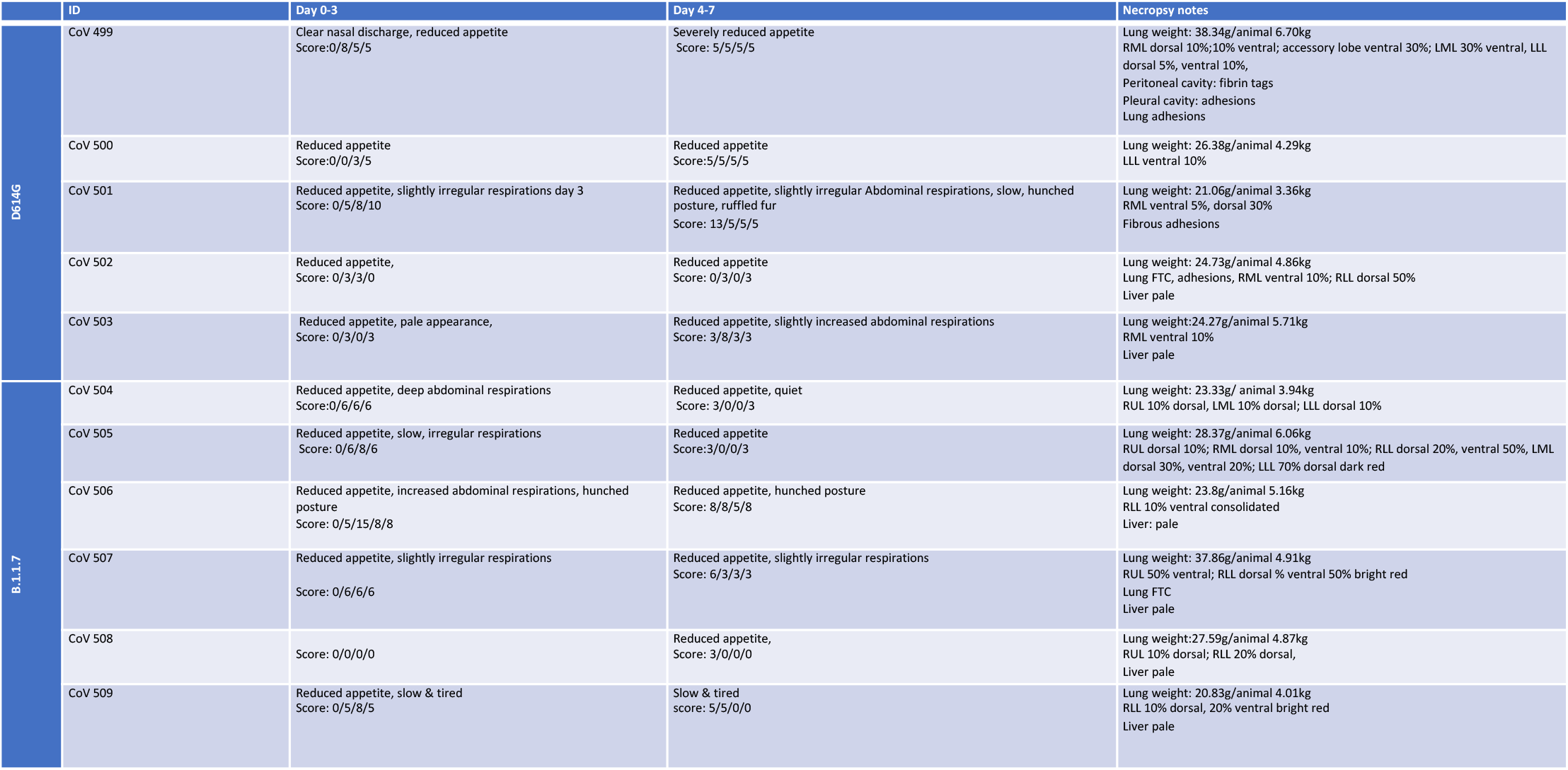
Clinical scoring and necropsy notes of infected animals. AGMs were scored daily for clinical signs of disease including changes in general appearance, respiration, food intake, fecal output as well as locomotion. Macroscopic scoring of organs was performed during necropsies (day 7 post-infection).

**Table S2:**
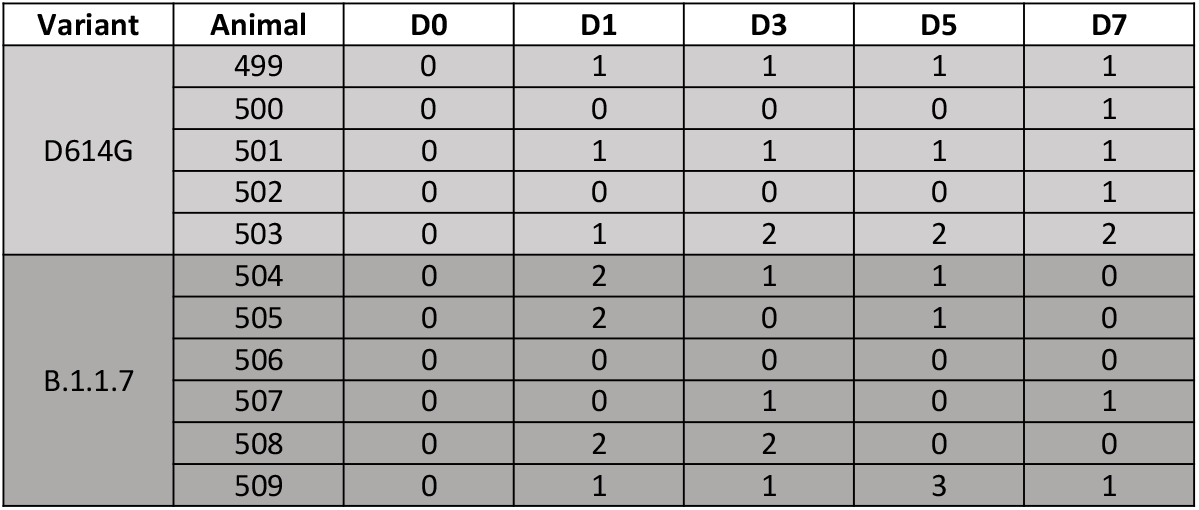
Radiographic scoring of lungs following infection. Ventro-dorsal and right/left lateral radiographs were taken on clinical exam days prior to any other procedures (e.g. bronchoalveolar lavage, nasal flush). Radiographs were evaluated and scored for the presence of pulmonary infiltrates by two board-certified clinical veterinarians according to a standard scoring system (*7*). Briefly, each lung lobe (upper left, middle left, lower left, upper right, middle right, lower right) was scored individually based on the following criteria: 0 = normal examination; 1 = mild interstitial pulmonary infiltrates; 2 = moderate interstitial pulmonary infiltrates, perhaps with partial cardiac border effacement and small areas of pulmonary consolidation (alveolar patterns and air bronchograms); and 3 = pulmonary consolidation as the primary lung pathology, seen as a progression from grade 2 lung pathology. At study completion, thoracic radiograph findings were reported as a single radiograph score for each animal on each exam day. To obtain this score, the scores assigned to each of the six lung lobes were added together and recorded as the radiograph score for each animal on each exam day. Scores can range from 0 to 18 for each animal on each exam day.

| Variant | Animal | D0 | D1 | D3 | D5 | D7 |
| --- | --- | --- | --- | --- | --- | --- |
| D614G | 499 | 0 | 1 | 1 | 1 | 1 |
|  | 500 | 0 | 0 | 0 | 0 | 1 |
|  | 501 | 0 | 1 | 1 | 1 | 1 |
|  | 502 | 0 | 0 | 0 | 0 | 1 |
|  | 503 | 0 | 1 | 2 | 2 | 2 |
| B.1.1.7 | 504 | 0 | 2 | 1 | 1 | 0 |
|  | 505 | 0 | 2 | 0 | 1 | 0 |
|  | 506 | 0 | 0 | 0 | 0 | 0 |
|  | 507 | 0 | 0 | 1 | 0 | 1 |
|  | 508 | 0 | 2 | 2 | 0 | 0 |
|  | 509 | 0 | 1 | 1 | 3 | 1 |

